# When population growth intensifies intergroup competition, female colobus monkeys free-ride less

**DOI:** 10.1101/2023.05.05.539387

**Authors:** T. Jean Arseneau-Robar, Julie A. Teichroeb, Andrew J. Macintosh, Tania L. Saj, Emily Glotfelty, Sarah Lucci, Pascale Sicotte, Eva C. Wikberg

## Abstract

In many social species, intergroup aggression is a cooperative activity that produces public goods such as a safe and stable social environment and a home range containing the resources required to survive and reproduce. In this study, we investigate temporal variation in intergroup aggression in a growing population of colobus monkeys to ask a novel question: “Who stepped-up to produce these public goods when the competitive landscape changed?”. Both whole-group encounters and male incursions occurred more frequently as the population grew. Males and females were both more likely to participate in whole-group encounters when monopolizable food resources were available, indicating both sexes engaged in food defence. However, only females increasingly did so over time, suggesting that when intergroup competition intensified, it was females who increasingly invested in home range defence. Females were also more active in male incursions at high population densities, suggesting they also worked harder to maintain a safe and stable social environment over time. This is not to say that males were chronic free-riders when it came to maintaining public goods. Males consistently participated in the majority of intergroup interactions throughout the study period, indicating they may have lacked the capacity to invest more time and effort.

## Introduction

Humans have modified the environments we inhabit to an unprecedented extent. We have drastically altered landscapes [1], hunted some species to extinction and enabled other to proliferate [2], and driven climate change on a global scale [3]. Consequently, animal populations have experienced dramatic changes in their risk of mortality, and the quality of the habitat they live in. For social species, the resulting changes in population size, food availability, and group composition impact the intensity of intergroup competition experienced [4,5]. However, we know very little about how group-living individuals behaviourally adapt to such changes in their competitive landscape.

During each encounter with a neighbouring group, every group member must weigh the costs versus benefits of participation [6,7] when deciding whether to engage in intergroup aggression. Participants pay an opportunity cost to fight [8], and can risk wounds, injuries, or even death [9–13]. But there are numerous potential benefits that could incentivize individuals to take this risk, such as maintaining or improving access to important resources like space, mates, food, water, or shelter [e.g., 6,7,14,15]. However, individuals may also participate for other reasons like defending themselves or their offspring [16–18], because they have been forcefully recruited into the fight [19], or to obtain a return benefit in the future. For example, they may support group members in the fight with the expectation of a reciprocal investment at a later time [20], or use intergroup aggression as a costly and honest signal of their genetic quality so to attract more mates [21]. Lastly, intergroup aggression may function to discourage or prevent extra-group individuals from trying to immigrate into the group [22,23]. Irrespective of the benefit(s) that motivate participation, intergroup aggression often produces public goods as a by-product.

Public goods are goods that are non-excludable, meaning that all group members can utilize them, regardless of whether they contributed to their production [24]. We posit that there are two public goods that intergroup aggression commonly produces: access to a home range containing the resources required to survive and reproduce, and a safe and stable social environment. In territorial species, intergroup aggression can function to delineate territory boundaries [13,25]. However, even in non-territorial species, the group that wins typically displaces the group that loses [26] and groups that consistently win intergroup conflicts can annex space from their neighbours [27,28]. Consequently, participation in intergroup conflicts can impact the quantity and quality of resources a group has access to within their home range or territory. Participation in intergroup conflicts can also prevent the forceful immigration of extra-group members [29–32]. When participants fail to repel unwanted immigrants, resident group members may be overthrown and/or evicted [22], infanticidal attacks may occur, and group members may emigrate to avoid the period of violence and instability that can follow [33–36]. Given the social upheaval that can result from such a takeover, intergroup aggression that repels potential immigrants can provide a safe and stable social environment for group members.

Public goods are vulnerable to collective action problems because individuals that pay the costs of producing the public good obtain lower net benefits than free-riding group members [24]. These payoff differences disincentivize participation, and can lead to a ‘tragedy of the commons’ in which no one produces the public good [24]. The ubiquity of intergroup conflicts across social species suggests that most social animals overcome collective action problems, at least to some extent.

When the public good(s) produced from intergroup aggression are viewed within the framework of the Volunteer’s Dilemma, the production curve is modelled as a step function [37,38]. The implication of this is that a threshold number of ‘volunteers’ are required to produce the public good. For example, a group may need to mobilize more active participants than the opposing group to win the fight [19,39]. In the Volunteer’s Dilemma, the payoffs are such that it is less costly to volunteer to produce the public good than it is to live without it [37,38]. For example, it is better to fight to maintain a home range than it is to live without consistent access to the resources found within it. When viewing participation in intergroup conflicts as a Volunteer’s Dilemma, the most relevant question to understanding public goods production is ‘Which group members volunteer, and in what conditions do they do so?’. Work to date has highlighted that it is often privileged individuals that have priority-of-access to the public good that are most likely to produce it [39,40]. For example, individuals who are high-ranking may obtain a disproportionate share of the resources within their home range, and so have the most to gain from volunteering. Quantitative studies support this supposition, as high-ranking individuals have consistently been found to participate in intergroup conflicts more frequently than low-ranking group members [e.g., 7,17,39,41–46].

Although we often talk about public goods ‘production’, it is notable that both of the public goods that arise from intergroup aggression need to be ‘maintained’ over time. Social groups must repeatedly engage in intergroup conflicts to maintain home range boundaries, and repeatedly repel intruders to maintain a safe and stable social environment. If changing social (e.g., population density) or ecological conditions (e.g., resource availability) alter the intensity of intergroup competition [4,5], more volunteers may be needed to win a fight and/or volunteers may need to engage in intergroup aggression more frequently to maintain these public goods. However, we are aware of no studies to date that have investigated temporal variation in volunteering to ask, “Who volunteers when the public good becomes harder to maintain?”. In this study, we examine participation in intergroup conflicts in a growing population of ursine (‘white-thighed’) colobus monkeys (*Colobus vellerosus*) to address this knowledge gap.

We examine contributions to two different public goods (home range defence and a safe and stable social environment) by investigating participation in two types of intergroup conflicts. Whole-group conflicts occur when two groups are in close proximity and one or more individuals from at least one group exhibits intergroup aggression. Because entire groups are involved, the outcome of these encounters likely determines who wins access to the resources currently contested, but could also impact home-range boundaries over the long-term. Thus, participation in whole-group conflicts likely produces the public good of a successfully defended home range and access to the resources within it. Incursions are when one individual, or a coalition of same-sexed individuals, approach another group without the bulk of their social group. Although incursions can be neutral/affiliative (i.e., prospecting) [29,31,46–48], they can also be highly aggressive, with one or more individuals attempting to forcefully immigrate into a group [29–32,49]. In ursine colobus, male incursions can result in take-overs, in which the resident alpha male is evicted, infanticidal attacks occur, and female group members emigrate to avoid the period of instability that can follow [33,34]. Thus, participation during male incursions likely produces the public good of a safe and stable social environment.

We hypothesized that the intensity of intergroup competition would increase as the population grew (i.e., increased in density), and that both whole-group conflicts and male incursions would increase in frequency over time as a result. We investigate how males and females each modified their propensity to participate in intergroup conflicts (i.e., volunteer) when it became more difficult to maintain public goods. Because female participation in intergroup conflicts often functions as food defence [7,42,46,50–53], we expected that females would be most likely to participate in whole-group conflicts when high-quality, monopolizable food resources (young leaves, fruits, seeds and flowers) were available within their home range, and that they would increasingly participate in food defence as the population size increased. Although male participation in intergroup conflicts is thought to primarily function as mate defence [15,54–58], studies on other species of black-and-white colobus have also found evidence for male food defence [59,60]. Therefore, we also expected males to participate when monopolizable food resources were at stake, and to increasingly do so as intergroup competition intensified. We further expected males to be the primary participants during male incursions, and to increasingly participate in these as the population grew and competition for mates intensified. From these predicted patterns of participation, we hypothesize that male volunteers would contribute to the production of both public goods (i.e., home range defence and a safe and stable social environment), whereas females would primarily volunteer to produce the public good of home range defence.

## Methods

### Study population

The study population of ursine colobus monkeys inhabits a 1.92 km^2^ semi-deciduous dry forest by the villages of Boabeng and Fiema in central Ghana (70 43’ N and 10 42’ W; ESM Fig. 1). The colobus monkeys at this site were traditionally protected by religious taboos. After a period during which the taboos eroded and only a few dozen monkeys remained, the elders approached Ghana Wildlife Division for governmental protection. They also started an ecotourism project – the Boabeng-Fiema Monkey Sanctuary – that became one of Ghana’s top tourist destinations [61,62]. As a result, the colobus population at Boabeng-Fiema has steadily grown over the past 40 years, increasing from a few dozen individuals to more than 300 by 2014 (ESM Fig. 2) [63–65]. We used data collected between 2000 and 2009, during which time the population increased from approximately 200 to 300 individuals.

**Figure 1.**
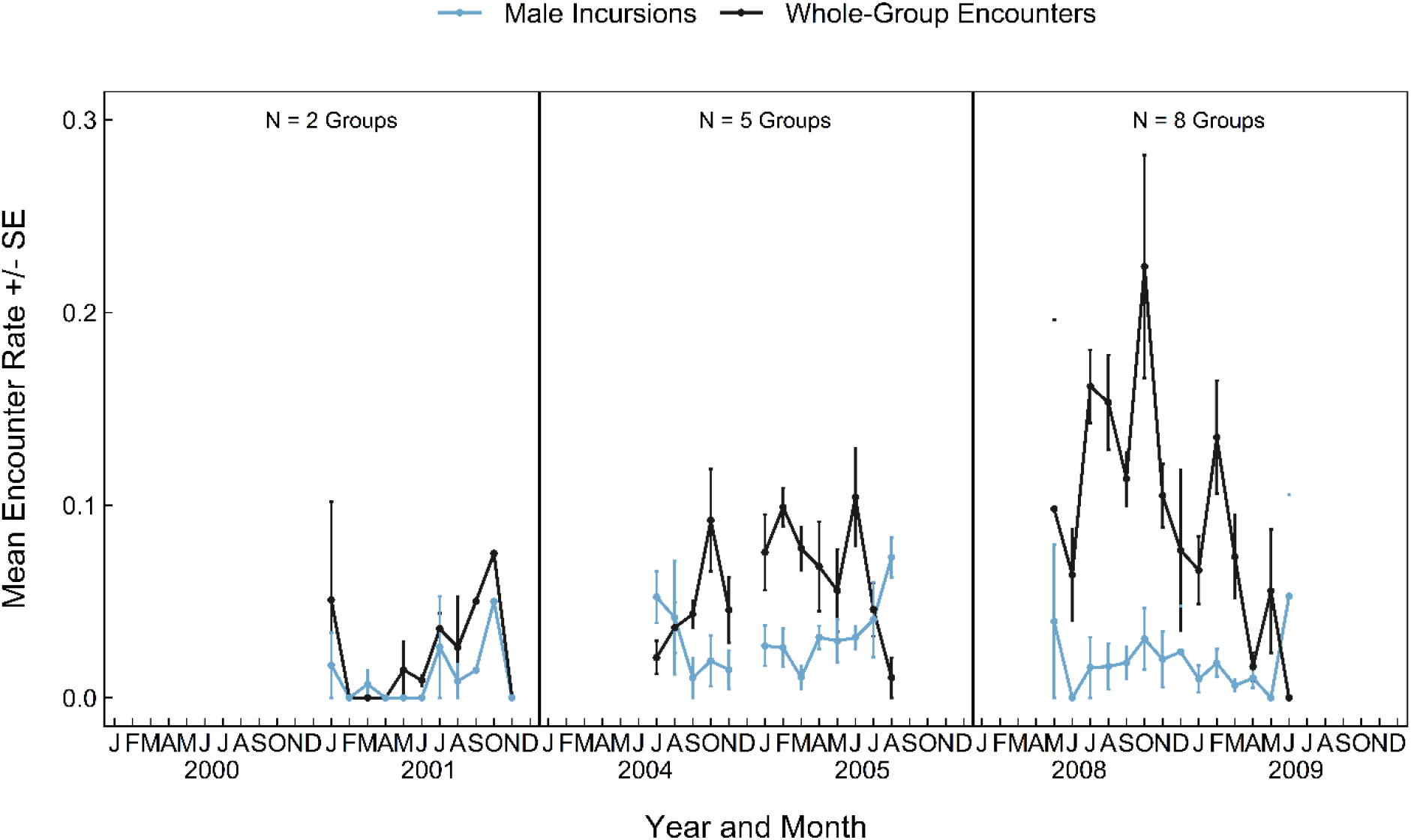
Mean rates of whole-group encounters and male incursions in ursine colobus monkeys, between 2001 and 2009. Mean values are the average monthly values among all study groups, and error bars represent standard errors.

**Figure 2.**
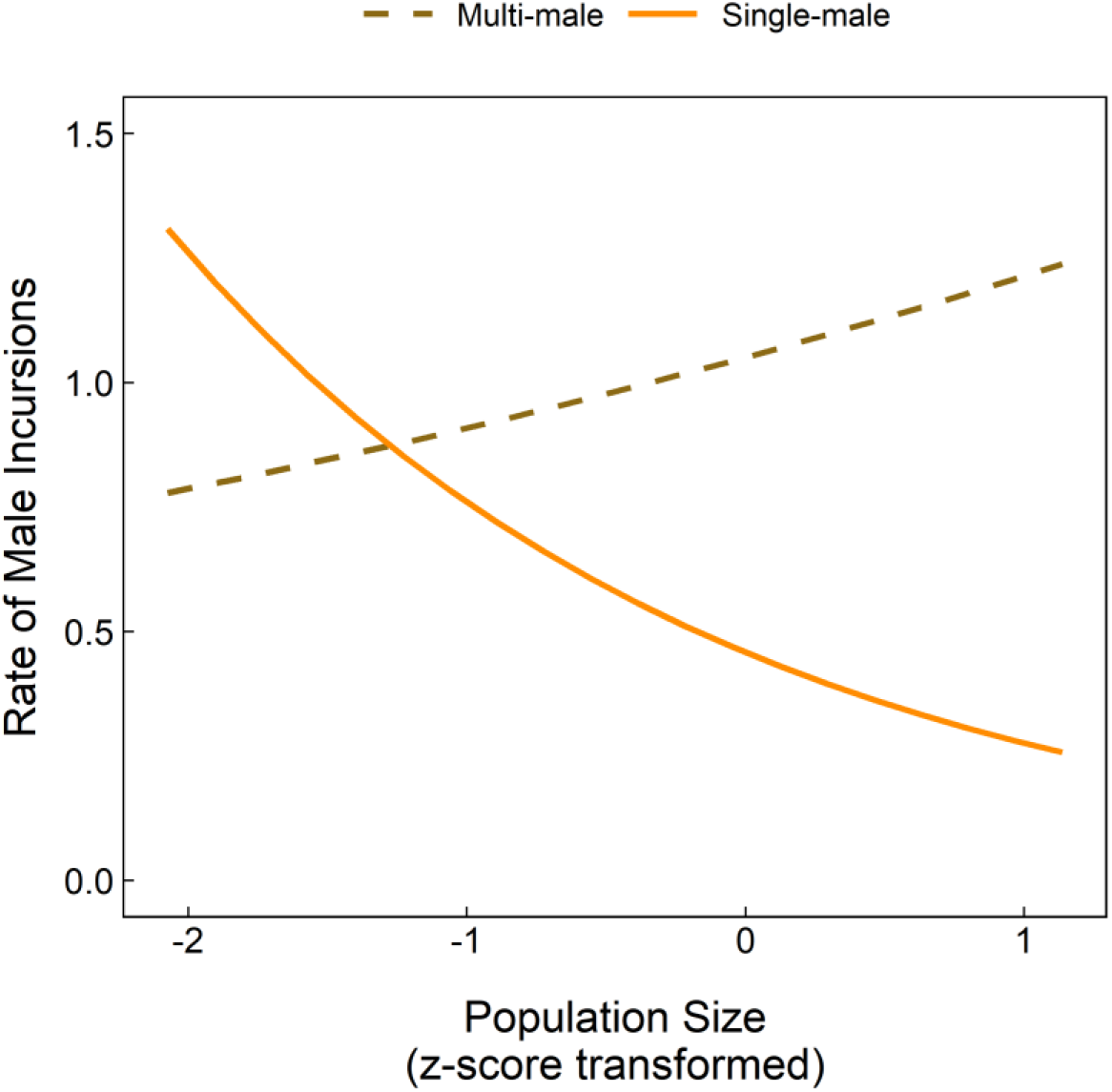
Predicted male incursion rate as a function of the interaction between population size and whether the group contained a single versus multiple adult males. Expected proportions were obtained by plotting the predictions of the GLMMs (Table 2), setting all additional predictors to their mean values (or median values for categorical variables). Measures of population size have been z-score transformed.

### Data collection

We utilized data collected by four different researchers between in three separate study periods (2000-2001, 2004-2005 and 2008-2009). To ensure consistency in behavioural data collection, each researcher overlapped with the next for at least one month in the field. Researchers collected data on group composition, the movements of the study groups, their diet, the distribution of important food trees, their monthly phenology, temperature and rainfall. Researchers also recorded the location of the centre-of-mass of the group every 30 minutes, and these locations were used to estimate annual home ranges for each study group, each study period. Researchers also collected aggressive participation *ad libitum* during two types of intergroup encounters: whole-group encounters, and incursions by one or more males. Two groups were deemed to be having a whole-group encounter when located within 50m of each other [58,66], and in conflict when aggressive behaviours, which included threat displays, chasing, and contact aggression [58,66], were observed. Incursions occurred when one individual, or a small number of same-sexed individuals, left their group (if they were from a bi-sexual group) and approached another group by themselves (i.e., their group was >50m away). The majority of incursions observed were by one or more males. Female incursions were observed but were at too low of a frequency to be analysed here.

### Data analyses

Because not all group members were individually recognized in the early study years, data on aggressive participation during intergroup encounters was summarized at the level of the sexes. These data ware also aggregated on a monthly basis to enable us to calculate encounter rates. We built two generalized linear mixed models (GLMMs) to investigate the factors that influenced the frequency of whole-group encounters, and the frequency of male incursions. The response variable in each model was the number of each type of encounter that each group experienced in a given month. We set the error structure to “Poisson”, used a “log” link function, and also included the observation hours for that group, that month, as an offset to control for differences in sampling effort. Study group was included as a random effect to account for the repeated sampling of groups over time [67]. We then built four separate GLMMs to examine the factors that drove both male and female participation during whole-group encounters and male incursions. The response variable in each male participation model was the number of encounters that at least one male group member participated aggressively in that month. Similarly, the response variable in each female participation model was the number of encounters that at least one female group member participated aggressively in that month. We set the error structure to “binomial” and used a “logit” link function. We also included the number of encounters each group experienced that month as a weight in each GLMM, and included study group as a random effect [67]. We did not include random slopes as our dataset did not support the more complex model structure [68–70]. Predictor variables in all models included estimated population size, monthly rainfall, an index of food availability within the group’s home range, and variables quantifying the numbers of adult males and females in the group.

Because the main goals of this study were to (1) determine whether the intensity of intergroup competition increased with population size, and (2) examine how males and females each responded to this, we included population size as a predictor variable in each of our GLMMs. Although the population of ursine colobus living at Boabeng-Fiema have been censused frequently since 1990, censuses were not conducted in every year that researchers were collecting data on intergroup encounters. Therefore, we used the censuses that were completed to create a linear model between year and population size, and then used the estimated population size from this model in our analyses (ESM Fig. 2).

The study site experiences a marked dry season (typically November to March) and two peaks in rainfall (May to July and September to October) (ESM Fig. 3a). This seasonal rainfall pattern leads to a peak of young leaves, seeds, fruits, and flowers in the dry season (ESM Fig. 3b) [71]. The plant part availability is reflected in the colobus diet consisting mostly of mature leaves in the rainy season and more high-quality food items in the dry season [71]. These high-quality food items, which are also those that tend to be more patchily distributed [72], and thus, are expected to increase the intensity of intergroup contest competition during the dry season [73,74]. Contrastingly, there is also evidence that groups may increase day ranges during the rainy season, which could lead to a higher number of intergroup encounters [75]. To control for these potential seasonal effects on the frequency of intergroup encounters and/or the participation of individuals in these encounters, we include the long-term average rainfall per calendar month (ESM Fig. 3a) as a predictor variable in all our models. We used long-term average rainfall per calendar month rather than the observed amount of rainfall for each month because rainfall data was not collected throughout the entire study period, but rather in 2001 and from 2003-2006.

**Figure 3.**
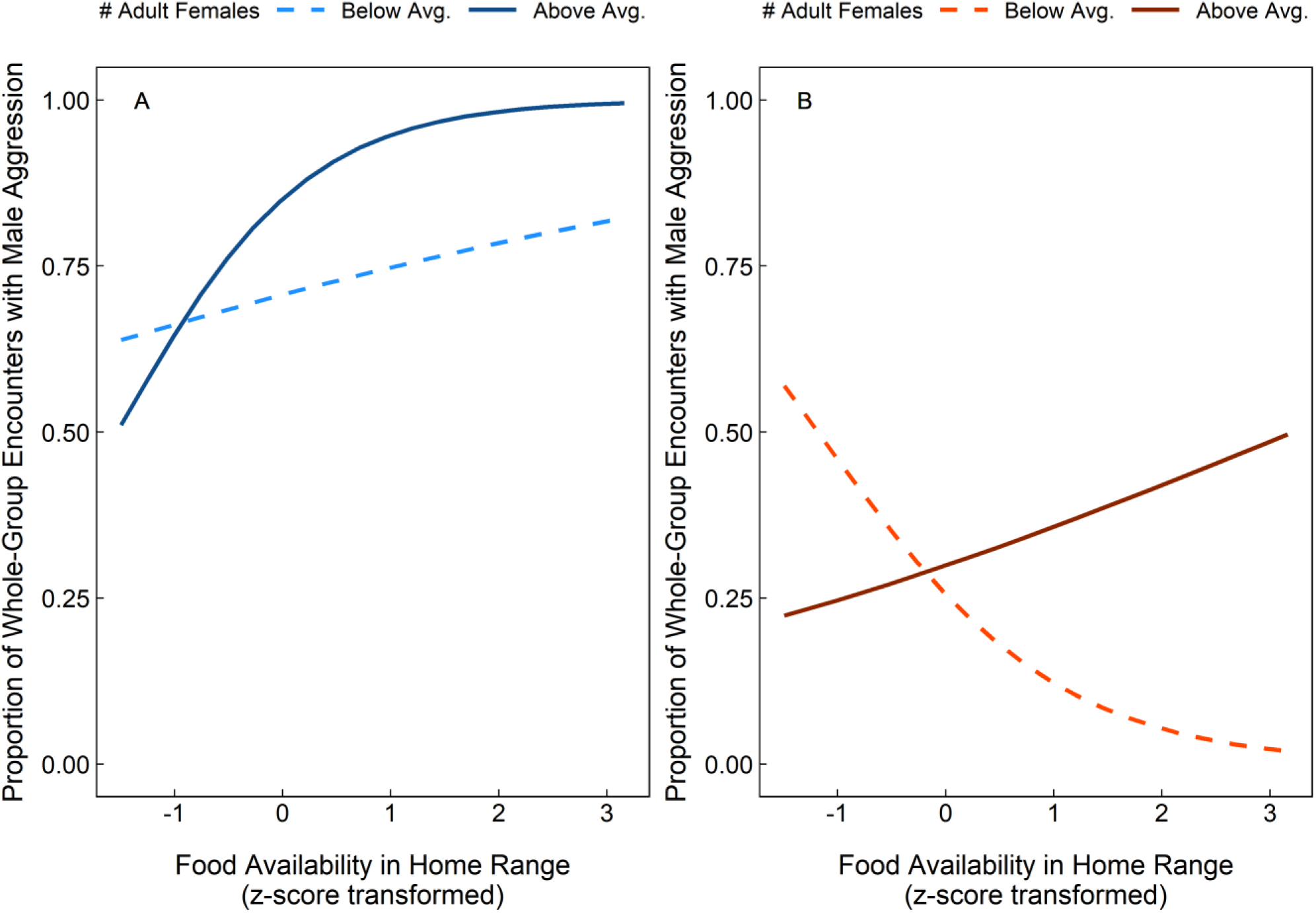
Predicted proportion of intergroup encounters in which (A) male and (B) female group members are expected to participate aggressively as a function of the interaction between the number of adult females in the group and the availability of young leaves, flowers, fruits and seeds within their home range. Expected proportions were obtained by plotting the predictions of the GLMMs (Table 1), setting all additional predictors to their mean values (or median values for categorical variables), and the number of adult females in the group to +1 SD above the mean (i.e., above average number of adult females in group) versus -1 SD below the mean (i.e., below average number of adult females). Measures of food availability have been z-score transformed.

Although the typical amount of rainfall each month should explain some of the variation in food availability (i.e., more young leaves, flowers, fruits and seeds available in the dry season), we also observed considerable variation in food availability among years, and among the home ranges of each of the study groups. Therefore, we also included a more precise estimate of current food availability for each study group as a predictor variable in our analyses. More specifically, we created an aggregated food-availability index (FAI) [76] that approximated the availability of monopolizable foods in the home range of each group, during each study period. This was accomplished by estimating each group’s home range boundaries, and then using data on the distribution of tree species important in their diet and their monthly phenology to calculate the monthly FAI for flowerbuds, flowers, unripe fruits, ripe fruits, unripe seedpods, ripe seedpods and young leaves (see Electronic Supplementary Materials for additional detail).

We included the number of adult females in each group as predictor variables in all our GLMMs, but changed the way we quantified male group composition. We used the number of adult males in our GLMMs examining rates because here we were primarily interested in how the number of males and/or females, and therefore the level of intragroup competition, influenced the intensity of intergroup contest competition. Conversely, in our GLMMs examining participation we used single-male versus multi-male as a predictor variable because previous work has shown differences in intergroup conflict outcome between single-male and multi-male groups [75], suggesting participation may also differ between these two types of groups. Six of the eight study groups were single male groups for at least part of the study period, and we observed groups to be single male across 40% of all group-months in the study period.

All predictor variables were *z*-score transformed prior to analyses [77] and we used pair-wise correlation coefficients and variance inflation factors [‘car’ package, version 3.0-7, 78] to ensure there was no multicollinearity among predictors (i.e., all VIF were < 3.0 and correlation coefficients < 0.8)[67,77]. We built all GLMM models using the ‘lme4’ package [version 1.1-21, 79] in R [80]. The ‘DHARMa’ package [81] was used to ensure that each GLMM did not suffer from over-or under-dispersion. We used the “confint.merMod” function in the ‘lme4’ package [79] to derive profile confidence intervals for each model parameter, and used these confidence intervals as well as likelihood ratio tests (comparing the full model to the model with each term removed) to assess the significance of each predictor variable and interaction term. We based our inferences on full models plus important interaction effects rather than using a stepwise procedure [82,83], and we did not interpret main effects if that predictor also featured in a significant interaction term. We also used a likelihood ratio test, in which we compared the full model to the null model (model including only the intercept, random effects, weight and offset (where applicable)) to assess the overall significance of each GLMM [67,84]. Lastly, the “r.squaredGLMM” function from the ‘MuMIn’ package [85] was used to obtain delta *R*^2^_GLMM(C)_ values for each model, as an estimate of model fit [86].

## Results

We observed 538 intergroup encounters (N = 409 whole-group encounters; N = 129 male incursions) during our 6306 observation hours and we were able to record participation data during 326 whole-group encounters and 127 male incursions. Males were equally as likely to participate aggressively in whole-group encounters (75% of N = 326) and male incursions (80% of N = 127) (Chi-squared test: *N* = 453, χ2 = 0.97, *p* = 0.325). Conversely, females were more likely to participate aggressively during whole-group encounters (35% of N = 376) than they were during male incursions (17% of N = 127) (Chi-squared test: *N* = 453, χ2 = 13.55, *p* < 0.001).

### Rates of intergroup encounters

The rate of whole-group encounters was strongly influenced by population size as these occurred at significantly higher frequencies over time (Fig. 1; Table 1). Groups that contained few adult females were also significantly more likely to experience whole-group encounters more frequently (Table 1). The effect of population size on the rate of male incursions was more complicated; multi-male groups were more likely to experience male incursions as population size increased, while single-male groups were less likely (Fig. 2; Table 2). Male incursions were also significantly less likely to occur when monopolizable food resources were abundant (Table 2), and a non-significant trend indicates there was a weak tendency for male incursions to occur more frequently in groups that contained few adult females (Table 2).

**Table 1.** Factors impacting the rate of whole-group encounters in ursine colobus monkeys, as well as the propensity of adult males and adult females to participate aggressively in these whole-group encounters.

| Model and Fixed Effects | *b*<br>Estimate | SE | $\chi^2$ | *p* | -95%<br>CI | +95%<br>CI |
| --- | --- | --- | --- | --- | --- | --- |
| **Whole-group encounter rate** |  |  |  |  |  |  |
| (Intercept) | -2.81 | 0.15 | - | - | - | - |
| Rainfall | -0.03 | 0.06 | 0.28 | 0.598 | -0.14 | 0.08 |
| **Population size** | **0.33** | **0.07** | **23.01** | **<0.001** | **0.19** | **0.48** |
| Food availability in home range | -0.07 | 0.07 | 0.89 | 0.346 | -0.22 | 0.07 |
| **Number adult females** | **-0.38** | **0.11** | **16.60** | **<0.001** | **-0.62** | **-0.18** |
| Number adult males | -0.10 | 0.07 | 2.09 | 0.148 | -0.24 | 0.03 |
| **Male aggression in whole-group encounters** |  |  |  |  |  |  |
| (Intercept) | 1.33 | 0.47 | - | - | - | - |
| Rainfall | 0.05 | 0.16 | 0.12 | 0.729 | -0.25 | 0.37 |
| **Population size** | **-0.42** | **0.22** | **3.85** | **0.049** | **-0.88** | **-0.00** |
| Food availability in home range | 0.67 | 0.25 | - | - | - | - |
| Number adult females | 0.43 | 0.32 | - | - | - | - |
| Single male vs multi-male | 0.10 | 0.48 | 0.05 | 0.830 | -0.84 | 1.10 |
| **Food availability \* Number adult females** | **0.47** | **0.22** | **4.75** | **0.029** | **0.05** | **0.90** |
| **Female aggression in whole-group encounters** |  |  |  |  |  |  |
| (Intercept) | -1.03 | 0.18 | - | - | - | - |
| **Rainfall** | **-0.44** | **0.15** | **9.44** | **0.002** | **-0.72** | **-0.16** |
| **Population size** | **1.08** | **0.19** | **38.37** | **<0.001** | **0.72** | **1.47** |
| Food availability in home range | -0.32 | 0.17 | - | - | - | - |
| Number adult females | 0.11 | 0.22 | - | - | - | - |
| Single male vs multi-male | 0.07 | 0.30 | 0.05 | 0.831 | -0.57 | 0.66 |
| **Food availability \* Number adult females** | **0.58** | **0.25** | **6.06** | **0.013** | **0.12** | **1.09** |

The whole-group encounter rate model explained half of the variation in IGE rates ( $R^2_{\text{GLMM(C)}} = 0.56$ ) and performed significantly better than the null model, which contained the intercept, offset, and random effects (likelihood ratio test: $N = 146$ , $\chi^2 = 53.35$ , $p < 0.001$ ). The models examining male participation (likelihood ratio test: $N = 146$ , $\chi^2 = 15.27$ , $p = 0.018$ ; $R^2_{\text{GLMM(C)}} = 0.55$ ) and female participation in whole-group encounters (likelihood ratio test: $N = 146$ , $\chi^2 = 72.02$ , $p < 0.001$ ; $R^2_{\text{GLMM(C)}} = 0.50$ ), both performed significantly better than the null models (i.e., models with intercept and random effects only). Significant predictors are presented in bold and trends are italicized.

**Table 2.** Factors impacting the rate of male incursions in ursine colobus monkeys, as well as the propensity of adult males and adult females to participate aggressively in these incursions.

| Model and Fixed Effects | *b* Estimate | SE | $\chi^2$ | *p* | -95% CI | +95% CI |
| --- | --- | --- | --- | --- | --- | --- |
| **Male incursion rate** |  |  |  |  |  |  |
| (Intercept) | -3.72 | 0.18 | - | - | - | - |
| Rainfall | -0.02 | 0.11 | 0.04 | 0.840 | -0.23 | 0.19 |
| Population size | 0.14 | 0.11 | - | - | - | - |
| **Food availability in home range** | **-0.31** | **0.16** | **5.20** | **0.023** | **-0.64** | **-0.04** |
| *Number adult females* | *-0.22* | *0.13* | *3.46* | *0.063* | *-0.56* | *0.01* |
| Single male vs multi-male | -0.83 | 0.29 | - | - | - | - |
| **Population size \* SM vs MM** | **-0.65** | **0.22** | **7.73** | **0.005** | **-1.08** | **-0.21** |
| **Male aggression in male incursions** |  |  |  |  |  |  |
| (Intercept) | 2.90 | 1.75 | - | - | - | - |
| *Rainfall* | *-0.61* | *0.35* | *3.08* | *0.079* | *-1.33* | *0.07* |
| Population size | -0.77 | 0.57 | 1.87 | 0.171 | -2.00 | 0.32 |
| **Food availability in home range** | **1.43** | **0.65** | **5.98** | **0.014** | **0.26** | **2.86** |
| **Number adult females** | **2.14** | **1.01** | **5.92** | **0.015** | **0.37** | **4.45** |
| Single male vs multi-male | -2.11 | 1.56 | 1.82 | 0.178 | -5.45 | 1.03 |
| **Female aggression in male incursions** |  |  |  |  |  |  |
| (Intercept) | -1.73 | 0.35 | - | - | - | - |
| Rainfall | -0.07 | 0.33 | 0.05 | 0.824 | -0.74 | 0.57 |
| **Population size** | **1.63** | **0.57** | **10.32** | **0.001** | **0.60** | **2.84** |
| Food availability in home range | 0.44 | 0.45 | 0.95 | 0.331 | -0.47 | 1.32 |
| Number adult females | 0.11 | 0.67 | 0.03 | 0.873 | -1.22 | 1.43 |
| *Single male vs multi-male* | *1.51* | *0.83* | *3.47* | *0.062* | *-0.08* | *3.32* |

The male incursion rate model explained a third of the variation in male incursion rates ( $R^2_{\text{GLMM}(C)} = 0.29$ ), and performed significantly better than the null model, which contained only the offset, intercept, and random effect (likelihood ratio test: $N = 146$ , $\chi^2 = 27.35$ , $p < 0.001$ ). The models examining male (likelihood ratio test: $N = 146$ , $\chi^2 = 19.73$ , $p = 0.001$ ; $R^2_{\text{GLMM}(C)} = 0.79$ ) and female participation in male incursions (likelihood ratio test: $N = 146$ , $\chi^2 = 30.77$ , $p < 0.001$ ; $R^2_{\text{GLMM}(C)} = 0.40$ ) both performed significantly better than the null models (i.e., models with the intercept and random effects only). Significant predictors are presented in bold and trends are italicized.

### Aggressive participation during whole-group encounters

Females were significantly more likely to participate aggressively in whole-group encounters at higher population densities (Table 1). Conversely, males showed the opposite pattern of behaviour as population size had a significant negative effect on male participation during whole-group encounters (Table 1). Females were also more likely to participate aggressively during whole-group encounters that occurred during the dry season (Table 1), which is the time of year that patchily distributed, monopolizable food resources were abundant (ESM Fig. 3). We also found that the interaction term between current food availability (i.e., the availability of young leaves, fruits, seeds and flowers) within the group’s home range and the number of adult females in the group to be statistically significant (Table 1). Plotting this interaction effect indicates that groups with many adult females, where intragroup feeding competition should be most intense, were more likely to participate aggressively when these contestable food resources were at stake (Fig. 3b). Conversely, in groups with few adult females, females were most active in the absence of food resources that are likely to induce contest competition (Fig. 3b). The interaction between current food availability and female group size was also a significant predictor of male participation during whole-group encounters (Table 1). Males were more likely to fight in whole-group encounters when patchily distributed, monopolizable food resources were abundant, but this propensity was stronger in groups that contained many adult females (Fig. 3a).

### Aggressive participation during male incursions

Males were most likely to participate aggressively during male incursions if their group contained many adult females (Table 2). Males were also more active during male incursions that took place when monopolizable food resources were abundant within their home range, and showed a weak tendency to participate in male incursions during the dry season when monopolizable food resources were available (Table 2). Notably, population size had no significant impact on male aggression during male incursions (Table 2). Conversely, females were significantly more likely to participate in repelling male intruders when living at a higher population density, as well as when they resided in a single-male group (Table 2).

## Discussion

Between the years 2000 and 2009, the population of ursine colobus monkeys living at Boabeng-Fiema grew by ∼150%, resulting in a significant increase in population density. We found that whole-group encounters became more frequent over time, as did the rate of male incursions, at least in multi-male groups. Thus, our findings indicate that as population density increased, so too did the intensity of intergroup contest competition. Patterns of participation in whole-group encounters suggest that the public good of a successfully defended home range may arise largely through male and female food defence. Both sexes were more likely to participate in whole-group encounters when monopolizable food resources were available within their home range and more likely to be depleted through scramble competition (i.e., when the number of females resident in the group was large). However, females were significantly more likely to engage in food defence as the population grew, whereas males were significantly less likely. Thus, it was females who increasingly volunteered when this public good became harder to maintain. Females were also the sex that increasingly invested in maintaining a safe and stable social environment, as their propensity to participate during male incursions correlated positively with population size. Conversely, male participation in male incursions was not related to population size, indicating that they consistently participated in ∼80% of male incursions. Thus, males may have hit their capacity to invest time and energy into public goods production, and so the increasing costs of maintaining these public goods in a changing competitive landscape largely fell to females.

### Aggressive participation in whole-group encounters

Our model of female participation in whole-group encounters showed that female ursine colobus monkeys were most active in whole-group encounters during the dry season, which is the time of year that high-quality, clumped food resources (i.e., young leaves, fruits, seeds and flowers) tended to be available. This particularly true for females living in groups with many adult females. Because female group size tends to correlate with total group size [87,88], groups with many adult females are likely to experience the highest levels of intragroup feeding competition, giving them the greatest incentive to fight for access to food resources. Female food defence has been found in the majority of studies that have examined female participation in intergroup conflicts [7,50–53] other resource that females have been seen to defend is space [7,46]. In this study, we found that females residing in groups with few adult females were most likely to participate in whole-group encounters when there were no monopolizable food resources at stake. This pattern could arise if females residing in smaller groups were primarily fighting to defend access to space. We observed that newly formed groups tended to be small compared to established groups. Given that the study site appears to be saturated with groups of colobus monkeys (ESM Fig. 1), these small new groups likely had to compete intensely with existing groups to establish and maintain a home range.

As seen in females, we also found evidence for food defence in males. Males were more likely to participate in both whole-group encounters and male incursions when monopolizable foods were available in their home range, and when their group contained many adult females (i.e., when intragroup scramble competition was likely to be high). That males show evidence for food defence during male incursions suggests that in some cases, incursions by a lone male or coalition of males are perceived as a whole-group encounter, even though the bulk of the group was >50 m away.

### Aggressive participation in male incursions

While male participation during male incursions was not impacted by population size, females were significantly more likely to contribute to repelling male intruders over time. Females presumably did so to mitigate the growing risk of infanticide associated with male takeovers [33]. Notably, in volunteering to maintain a safe and secure social environment, females also provided a mate-defence service to male group members as a by-product. Males able to maintain exclusive access to a group of females may have been more likely to obtain this service, as females showed a weak tendency to participate in male incursions when living in single-male groups. Females may have been more likely to assist in repelling extra-group males when residing in a single-male group, and facing a coalition of males, as the resident male would be at a numeric disadvantage in such cases. Evidence for such a strategy has been found in black howler monkeys, where male takeovers also carry the risk of infanticide. Here, playback studies have shown that females are most likely to participate in intergroup encounters when the males in their group are outnumbered [89].

## Conclusions

Many populations of social animals have likely experienced drastic changes in the intensity of intergroup competition they experience, either because they live in an area that has suffered habitat loss or degradation, and/or their population has increased. However, only a couple of studies have examined how such changes impact competitive dynamics between social groups [5,27], and none have examined how patterns of volunteering versus free-riding change in response to this evolving competitive landscape. Here, we use data collected on a growing population of ursine colobus monkeys to ask a novel question: when intergroup competition intensifies, such that it becomes more difficult to maintain home range boundaries and prevent immigration, who steps up and volunteers to produce these public goods. We found that it was females who increasingly fought for the resources that are limiting to their own fitness (i.e., food and space), and they also increased the time and effort invested in preventing male takeovers. This is not to say that males were free riders who exploited the efforts of their female group members. Males participated in the majority of intergroup encounters experienced, while females only participated in 35% of whole-group encounters and 17% of male incursions. Thus, while females were the sex who increasingly invested in home range defence and the maintenance of a safe and stable social environment when it became more difficult to maintain these public goods, this may have been because they were the sex with the greatest capacity to do so.

## Supporting information

Electronic Supplementary Materials

