## Supplementary material for "When population growth intensifies intergroup competition, female colobus monkeys free-ride less": Electronic Supplementary Materials

### Supplementary Methods

#### *Study population*

The study population of ursine black-and-white colobus monkeys (*Colobus vellerosus*) inhabits a 1.92 $km^2$ semi-deciduous dry forest by the villages of Boabeng and Fiema in central Ghana (70 43' N and 10 42' W; ESM Fig. 1). The colobus monkeys at this site were traditionally protected by religious taboos. After a period during which the taboos eroded and only a few dozen monkeys remained, the elders approached Ghana Wildlife Division for governmental protection. They also started an ecotourism project – the Boabeng-Fiema Monkey Sanctuary – that became one of Ghana's top tourist destinations [1,2]. As a result, the colobus population at Boabeng-Fiema has steadily grown over the past 40 years, increasing from a few dozen individuals to more than 300 by 2014 (ESM Fig. 2) [3–5]. This study utilizes data collected between 2000 and 2009, during which time the population increased from approximately 200 to 300 individuals.

#### *Estimating food availability for each group over time*

The study site experiences a marked dry season (typically November to March) and two peaks in rainfall (May to July and September to October) (ESM Fig. 3a). This seasonal rainfall pattern leads to a peak of young leaves, seeds, fruits, and flowers in the dry season (ESM Fig. 3b) [6]. The plant part availability is reflected in the colobus diet consisting mostly of mature leaves in the rainy season and more high-quality food items in the dry season [6]. These high-quality food items, which are also those that tend to be more patchily distributed [7,8], are expected to increase the intensity of intergroup contest competition during the dry season [9–12].

Although the typical amount of rainfall each month should explain some of the variation in food availability (i.e., more young leaves, flowers, fruits and seeds available in the dry season), we also observed considerable variation in food availability among years, and among the home ranges of each of the study groups. Therefore, we created an aggregated food-availability index (FAI) that approximated the availability of monopolizable foods in the home range of each group, during each study period. We used the 'adehabitatHR' package [13] to estimate the utilization distribution using the kernel density estimate approach [14], setting the smoothing parameter to the reference bandwidth. Home range boundaries were derived from the 95% contour of the utilization distribution. We used the observed diet of each study group [6,15, Wikberg unpublished data], during each study period, to create a list of 28 'important food-species' that comprised at least 5% of the annual diet of at least one group. We mapped the location of each individual tree, having a diameter at breast-height (DBH) of at least 40 cm, of these important food-species across the study area in 2000-2001 [6]. We used this detailed tree map to determine the total DBH for the important food-species within each annual home range. We used DBH because this metric correlates well with canopy size and overall tree productivity [16]. During our monthly phenology sampling, the abundance of flowerbuds and flowers, unripe and ripe fruits, unripe and ripe seedpods, and young and mature leaves were estimated for three to seven (mean = 5) representative trees of these important food species [17]. Observers gave each plant part a score from 0 to 4 (0 = none, 1 = 1% to 25%, 2 = 26% to 50%, 3 = 51% to 75%, 4 = 76% to 100% canopy cover) to index the canopy cover of that plant part; the maximum total score within each category (e.g., flowers, fruits, seeds, leaves) was 4 [6]. We calculated the average monthly phenology score for each plant part, of each species, and multiplied these average scores by the total DBH of that species in each home range, to get an index of food availability (FAI) for each plant part of each species, for each group [18,19]. We then created an aggregated FAI by summing the FAIs of flowerbuds, flowers, unripe fruits, ripe fruits, unripe seedpods, ripe seedpods and young leaves for all important food-species. Because these high-quality foods are also those that tended to have more patchy distributions, they are also expected to be those that would incite intergroup contest competition.

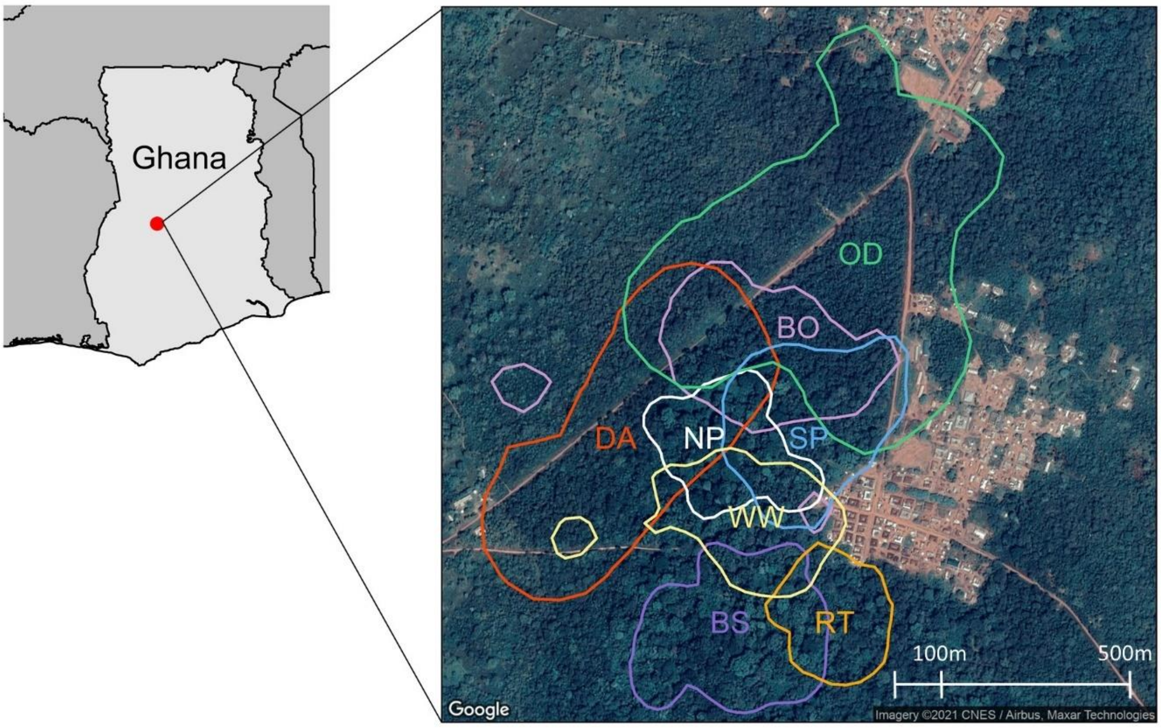


**ESM Figure 1.** Home ranges of the eight study groups of ursine colobus monkeys at Boabeng-Fiema Monkey Sanctuary in Ghana, during the 2008-2009 study period.

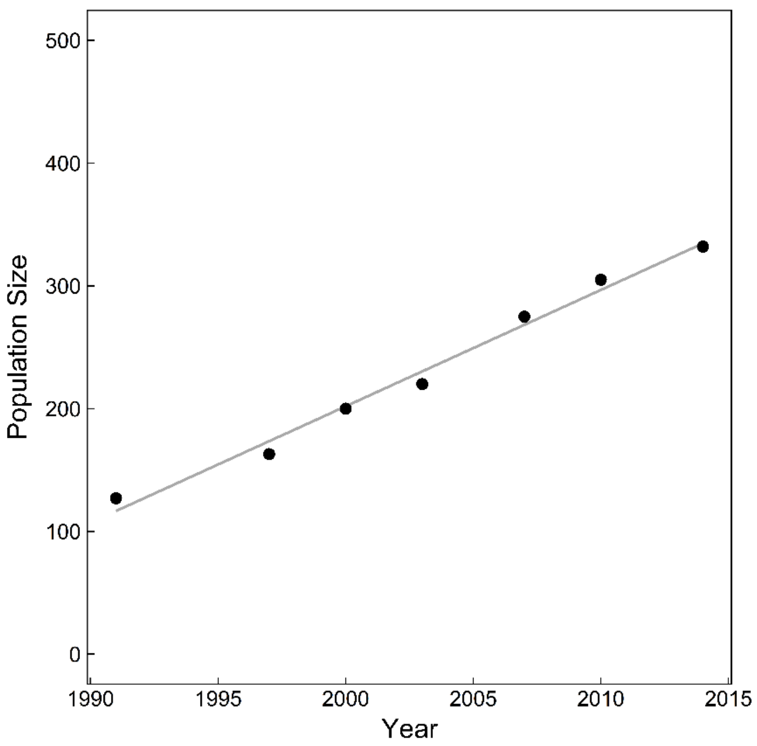


**ESM Figure 2.** Size of the population of ursine colobus monkeys living at the Boabeng-Fiema Monkey Sanctuary in Ghana between 2000 and 2014.

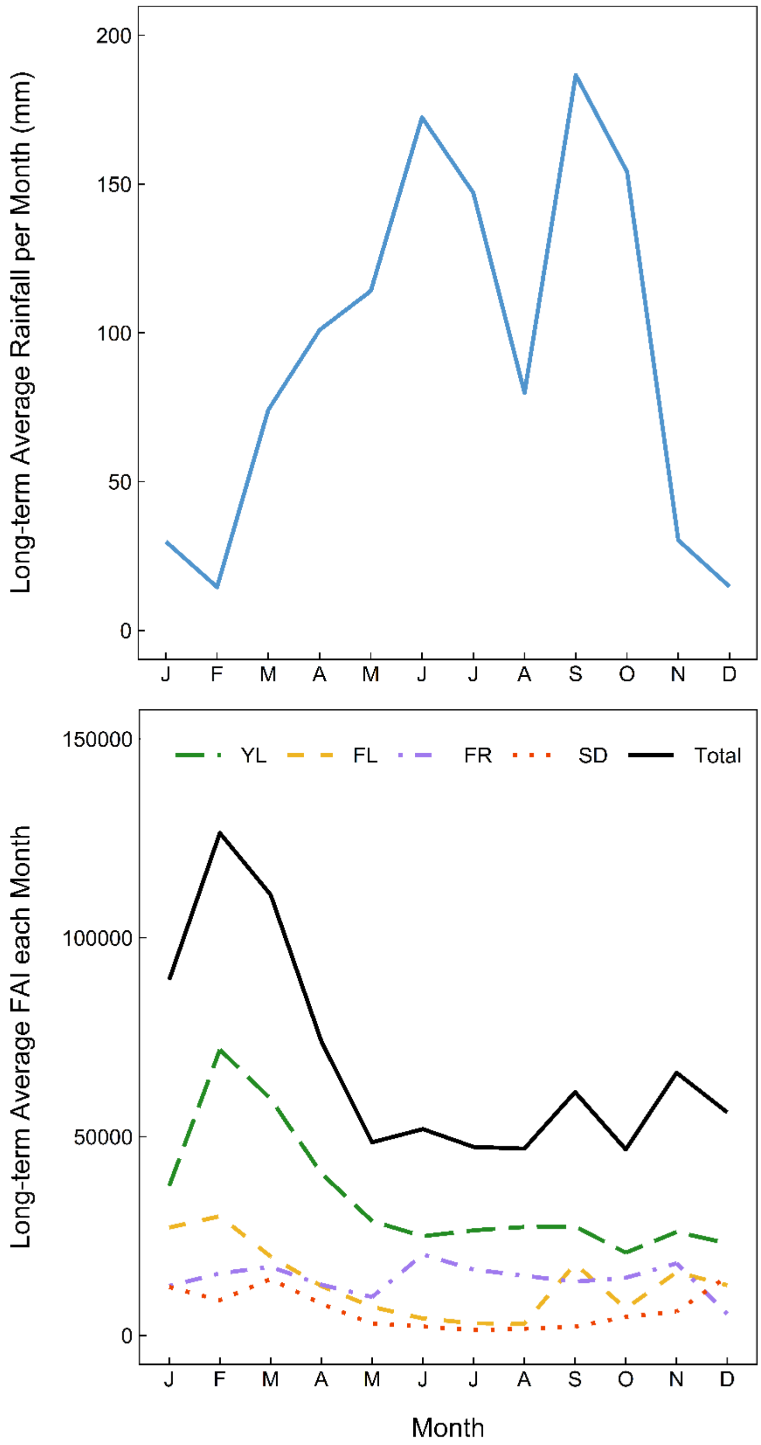


**ESM Figure 3.** Seasonal variation in rainfall and the availability of young leaves, flowers, fruits and seeds in the study area.
